# Inferring Single-Cell RNA Kinetics from Various Biological Priors

**DOI:** 10.1101/2024.05.21.595179

**Authors:** Yanshuo Chen, Zhengmian Hu, Ke Ni, Site Feng, Wei Chen, Heng Huang

## Abstract

In the context of transcriptional dynamics modeled by ordinary differential equations (ODEs), the RNA level in a single cell is controlled by specific RNA kinetics parameters, which include transcription rate, splicing rate, and degradation rate. Investigating these single-cell RNA kinetics rates is pivotal for understanding RNA metabolism and the heterogeneity of complex tissues. Although metabolic labeling is an effective method to estimate these kinetics rates experimentally, it is not suitable for current large-scale conventional single-cell RNA sequencing (scRNA-seq) data. Moreover, existing methods for scRNA-seq often either neglect certain specific kinetics parameters or use inappropriate ways to fit the parameters. To address these issues, we introduce scRNAkinetics, a parallelized method that fits the kinetics parameters of the ODE for each cell using pseudo-time derived from various biological priors (*e*.*g*. cell lineage tree and differentiation potential). This approach allows for the estimation of the relative kinetics of each cell and gene in a scRNA-seq dataset. Validated on simulated datasets, scRNAkinetics can accurately infer the kinetics rates of transcription boosting, multi-branch, and time-dependent RNA degradation systems. Nevertheless, the inferred kinetics trends are concordant with previous studies on metabolic labeling and conventional scRNA-seq datasets. Furthermore, we show that scRNAkinetics can provide valuable insights into different regulatory schemes and validate the coupling between transcription and splicing in RNA metabolism. The open-source implementation of scRNAkinetics is available at https://github.com/poseidonchan/scRNAkinetics.

## 1 Introduction

RNA metabolism, including the complex regulated processes of transcription, splicing, and degradation, is essential for cellular functionality. Dysfunctions in these processes are associated with a wide range of diseases, including cancer, [1,2,3] and neurological diseases [4]. High-throughput RNA sequencing, especially single-cell RNA sequencing (scRNA-seq) coupled with metabolic labeling, has emerged as a potent tool for studying these fundamental processes [5,6,7]. This approach allows for the quantitative investigation of genome-wide transcript expression levels and dynamics, providing insights into the complex nature of RNA regulation.

To analyze the dynamic relationship between the quantities of unspliced and spliced messenger RNA, a mathematical framework using a system of ordinary differential equations (ODEs) is employed to model the RNA dynamics [8]. In this set of ODEs, the changing rate of the unspliced transcript is determined by the transcription rate and the splicing of the current unspliced transcript, and the changing rate of the spliced transcript is affected by splicing and degradation only. Based on this model, many methods have been developed to infer the kinetic rates with various assumptions. Before single-cell resolution, methods like DTA [9], DRiLL [10], INSPEcT [11], and pulseR [12] were developed for bulk RNA sequencing with metabolic labeling data. For conventional scRNA-seq data, velocyto [13] and scVelo [14] can estimate gene-specific constant kinetic parameters while DeepVelo [15], cellDancer [16], and DeepKINET [17] can estimate cell- and gene-specific kinetic parameters based on the quantified unspliced and spliced RNA expression level [13,18]. However, beyond these methods’ capabilities, they also have their own limitations. For example, bulk RNA sequencing and metabolic labeling scRNA-seq are either not able to observe the nuance between different cells or need additional experimental costs, which limits their usage in the current era. For other methods developed for conventional scRNA-seq, the key problem is that simultaneously inferring time ordering and kinetic parameters is very challenging and almost impossible. Specifically, within the RNA dynamics model, the ODEs yield an analytic solution. This solution reveals that the abundance of spliced RNA is a function of both the abundance of unspliced RNA and the time point. However, this presents a challenge: the equation includes four unknowns – latent time and three kinetic parameters – yet provides only one equation to determine them. Therefore, velocyto and scVelo employ the steady-state assumption and treat kinetic parameters as constant among cells to infer them and the latent time. The recently developed cell- and gene-specific kinetics estimation methods use different neural networks to project RNA abundance to these kinetic parameters and then fit the neural network based on a possible future direction. Therefore, the efficacy of these methods is dependent on network initialization, and the process of fitting kinetic parameters is not rigorous compared to using the ODE solver. Among these methods, only cellDancer is able to infer the relative kinetic parameters for transcription, splicing, and degradation. In contrast, DeepVelo and DeepKINET are limited to inferring only the splicing and degradation rates.

To address these issues in estimating cell- and gene-specific kinetic parameters for conventional scRNA-seq data, it is necessary to first establish a correct cell order, assign a time point to each cell, and then infer the kinetic parameters, thus avoiding the dilemma of simultaneous inference of kinetics and time in the ODEs. In conventional scRNA-seq datasets, the real-time point is not available due to the destructive nature of the sequencing process. So we use the assumption that a cell’s near-future state is akin to one of the existing cells in the dataset, with a near-constant velocity during this transition, implying that the distance between these two cells correlates positively with the time interval of the transition. Then, instead of seeking the real-time, we can use the pseudo-time derived from various biological prior with various methods for kinetic parameters estimation. The first widely used prior is the stem cell marker prior shown in previous pseudo-time methods [19,20], in which the root cell can be determined by heuristic stem cell and special gene marker in a specific tissue. The second one is the differentiation potential prior. Previous works have shown that the protein-protein interaction networks or the number of expressed genes in a cell can be used to accurately estimate cell differentiation potential in many tissues [21,22]. The third one is the RNA velocity prior. This prior is shown in the original velocyto paper, where the researchers find that the RNA abundance in a developmental tissue usually experiences both transcription induction and repression stages [13]. The fourth prior is the cell lineage from single-cell lineage tracing data. Recently, PhyloVelo has been able to incorporate cell lineage information into the RNA velocity framework [23]. This is achieved by using monotonically expressed genes to accurately estimate the direction of cell differentiation. By integrating these priors, we can infer the correct direction of cell differentiation and pseudo-time for nearly all scRNA-seq datasets. However, the finer and more intriguing details, such as cell- and gene-specific kinetic parameters, remain largely unexplored.

In this light, we developed our method scRNAkinetics to infer the single-cell RNA kinetic parameters. scRNAkinetics assumes each cell and its neighbors have the same parameters, then the kinetic parameters are fitted by forcing the earliest cell to evolve into other later cells according to the ODEs (Fig.1). Of note, contrasting with methods like cellDancer that treat the system as underdetermined with the input being the abundance of two different stages of RNA and output as three kinetic parameters, our approach, which treats the kinetics parameters in a cell neighborhood as the same, transforms the underdetermined system into an overdetermined one, thereby enabling more reliable parameter estimation.

**Fig. 1.**
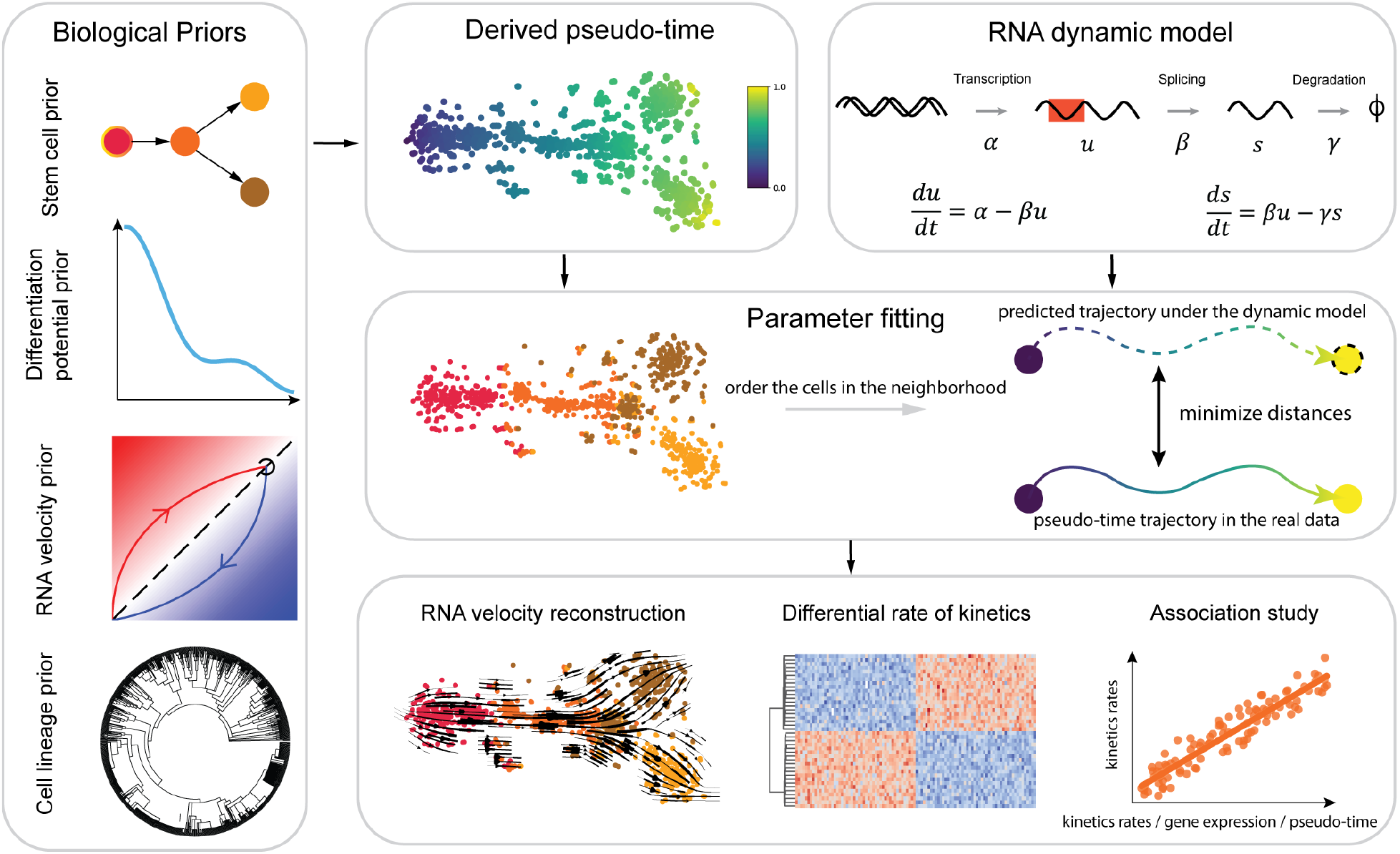
Overall illustration of scRNAkinetics workflow. scRNAkinetics leverages the pseudo-time trajectory derived from multiple biological priors combined with a specific RNA dynamic model to accurately infer the RNA kinetics for scRNA-seq datasets. scRNAkinetics assumes each cell and its neighborhood have the same kinetic parameters and fit the kinetic parameters by forcing the earliest cell evolve into later cells on the pseudo-time axis. After repeating this process for all cells in the dataset, it provides cell- and gene-specific kinetic parameters. These parameters are pivotal for conducting comprehensive bioinformatic analyses and offering insights into the detailed kinetics of gene expression at the single-cell level.

In this work, through the validation on both simulated and real datasets, we demonstrate scRNAkinetics can accurately estimate the kinetic parameters and handle complicated kinetics such as transcription boosting, multi-branch, and time-dependent kinetic parameter systems. Moreover, scRNAkinetics can lead to the gene regulation scheme findings that are congruent with previous metabolic labeling experiments [6]. Furthermore, we find a strong correlation between the estimated RNA transcription rate and splicing rate, aligning with previous studies that suggest a coupling between transcription and splicing processes in the cell nucleus [24].

## 2 Results

### 2.1 scRNAkinetics overview

As shown in Fig.1, our method is designed to take the pseudo-time derived from various biological priors together with a specific RNA dynamic model as input and then output estimated kinetic parameters for each cell and gene. The cell- and gene-specific kinetic parameters can be used to reconstruct velocity and other downstream analyses such as differentially expressed kinetics analysis between cell groups and association study. Hence, the first step is to derive an appropriate pseudo-time estimation based on some biological priors. In practice, the choice of prior is dependent on the data type and users’ professional knowledge. For example, if the sequencing data includes cell lineage information, users can employ PhyloVelo [23] to derive a reliable pseudo-time. If the data is conventional scRNA-seq, users can choose different tools based on their understanding of the sequenced tissue. When the stem cell type is not clear, they can use SCENT [21] to derive reliable pseudo-time. Alternatively, when the stem cell type is well known, they can use palantir [20] for this purpose. With a reliable pseudo-time estimation, scRNAkinetics can then estimate the parameters of the RNA dynamic model for each cell and gene in high-resolution mode. In this mode, it is assumed that each cell and its neighbors, defined by the expression distance in the single-cell analysis pipeline, share the same set of kinetic parameters. For the selected cell’s neighborhood, scRNAkinetics first selects the earliest cell in the neighborhood on the pseudo-time axis and subsequently predicts the future states of this cell under the RNA dynamic model at all future pseudo-time points. After having the predicted cell trajectory, we optimize the kinetic parameters by minimizing the distances between the predicted trajectory and the pseudo-time trajectory derived from the reliable biological prior. After that, we repeat this process for all cells and their neighbors to obtain the full kinetic parameters estimation of the whole dataset. Thus, the kinetic parameters would vary across cells and offer the ultimate resolution for us to investigate the kinetics variation in a tissue. With the estimated kinetic parameters, we can substitute them into the RNA dynamic model to obtain the unspliced and spliced RNA velocity and analyze the kinetic rates trends in a development tissue.

### 2.2 scRNAkinetics correctly resolve the kinetic rates on simulated datasets

In order to confirm that our method can be applied in principle, we evaluated our method on simulated data first, where the data was generated using the RNA dynamic model in four different scenarios: conventional system, transcription boosting system, multi-branch system, and time-dependent degradation rate system (see details of the simulation process in Methods). Of note, we started the evaluation without adding any noise to validate the feasibility of our method. As we can see, scRNAkinetics can automatically distinguish the ODEs system transition in the conventional and transcription boosting settings (Fig. 2a,b), while maintaining other kinetic parameters almost unchanged (Fig. 2a,b). Another challenging scenario is the multi-branch dynamic system, which usually impedes the correct RNA velocity estimation in many real-world datasets. Here the scRNAkinetics can also discern the transcription rate into two levels and identify other kinetic rates as unchanged (Fig. 2c). These three scenarios only include one-time kinetic rate change and thus may not be very practical in the real world. Following previous studies, where they have found the time-dependent rate in the development tissue [25,26], we simulated the time-dependent degradation rate and evaluated scRNAkinetics on it. The result shows that the estimated degradation rate is highly correlated with the ground truth (Fig. 2d).

**Fig. 2.**
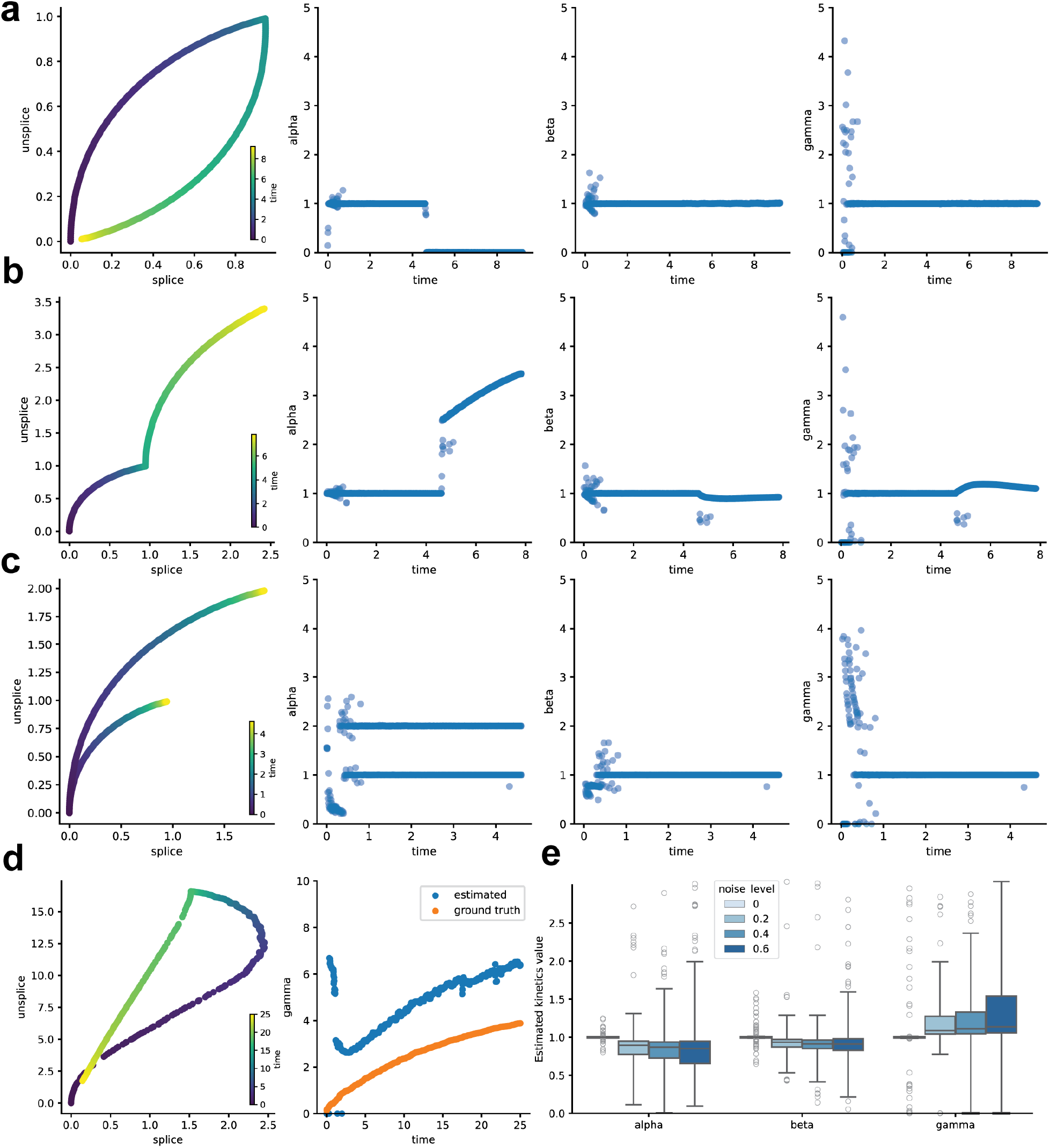
scRNAkinetics’ verification on synthetic data. **a**, Left most panel illustrates the simulated ODE trajectory of conventional dynamics. The color represents the specific time point of each data point on the plot. Three other panels are the estimated kinetic rate of transcription, splicing, and degradation plotted against time progression. **b**, Left most panel shows the simulated ODE trajectory of transcription boosting. Other elements’ meaning is the same as **a. c**, Left most panel shows the simulated ODE trajectory of multi-branch dynamics. Other elements’ meaning is the same as **a. d**, Left panel represents the simulated ODE trajectory of time-dependent kinetic rates. Right panel shows the time-dependent degradation rate. The ground truth is especially displayed since the estimation is not as perfect as previous ones due to the KNN smoothing process. We can still see the correlation between estimation and ground truth is very high. **e** Noise impacts the accuracy of scRNAkinetics. It demonstrates the impact of noise on the kinetic parameters estimation of the induction branch within the conventional scenario, depicting a clear trend where increased noise levels correspond to a larger variance in the rate estimations.

Although the estimation of scRNAkinetics is reasonable in most cases, we also find some potential caveats in the estimation process. The first one is the way to construct KNN in a dataset. Since we assume a cell and its neighborhood share the same kinetics, searching for a proper KNN would be crucial for the model performance. In the simulated datasets, especially the multi-branch scenario, it is hard to find a proper KNN that can distinguish different branches when the two branches are very close on the gene’s phase plot (Fig. 2c). However, we can hypothesize this issue could be mitigated when the KNN graph is built on more components (*e*.*g*. PCA space or full gene space) because different branches are expected to be distinguishable in high-dimensional space. Another issue that may lead to the inaccurate estimation is the KNN smoothing process since it averages the expression in the neighborhood and thus loses the original expression level. Hence, the estimation for the time-dependent kinetic rate is not as accurate as the constant kinetic rate but remains the high correlation between the ground truth (Fig. 2d).

Following the initial validation of scRNAkinetics, we then tested how the noise would affect the model estimation. We introduced varying degrees of Gaussian noise into the induction branch in the conventional system (*α* = 1, *β* = 1, *γ* = 1, see more details in Methods) and evaluated scRNAkinetics performance for all three parameters (Fig. 2e). The findings indicate an increase in estimation variance with escalating noise levels. However, the median estimation values consistently aligned closely with the true values, demonstrating the robustness of scRNAkinetics against noise interference.

### 2.3 scRNAkinetics is congruent with metabolic labeling study

To assess the scRNAkinetics performance on the real-world datasets, we introduced the metabolic labeling scEU-seq intestinal organoid dataset which can provide RNA synthesizing and degradation rate estimation [6]. This dataset includes several cell types which can be divided into three branches: the stem cell, the enterocytes, and the secretory cells (Fig 3a). The expected differentiation trajectory is that the stem cells are differentiated into two branches. To incorporate this prior, we employed the UnitVelo by setting the stem cells as root to estimate the latent time (Fig 3d) [27], then we used scRNAkinetics to estimate cell-specific kinetic parameters and reconstruct the RNA velocity (Fig 3b). We also compared it with other kinetics estimation methods, namely cellDancer, DeepVelo, and DeepKINET (Fig 3c,e,f). The results suggest only cellDancer can properly estimate the correct velocity direction while other methods fail and predict backward flows.

**Fig. 3.**
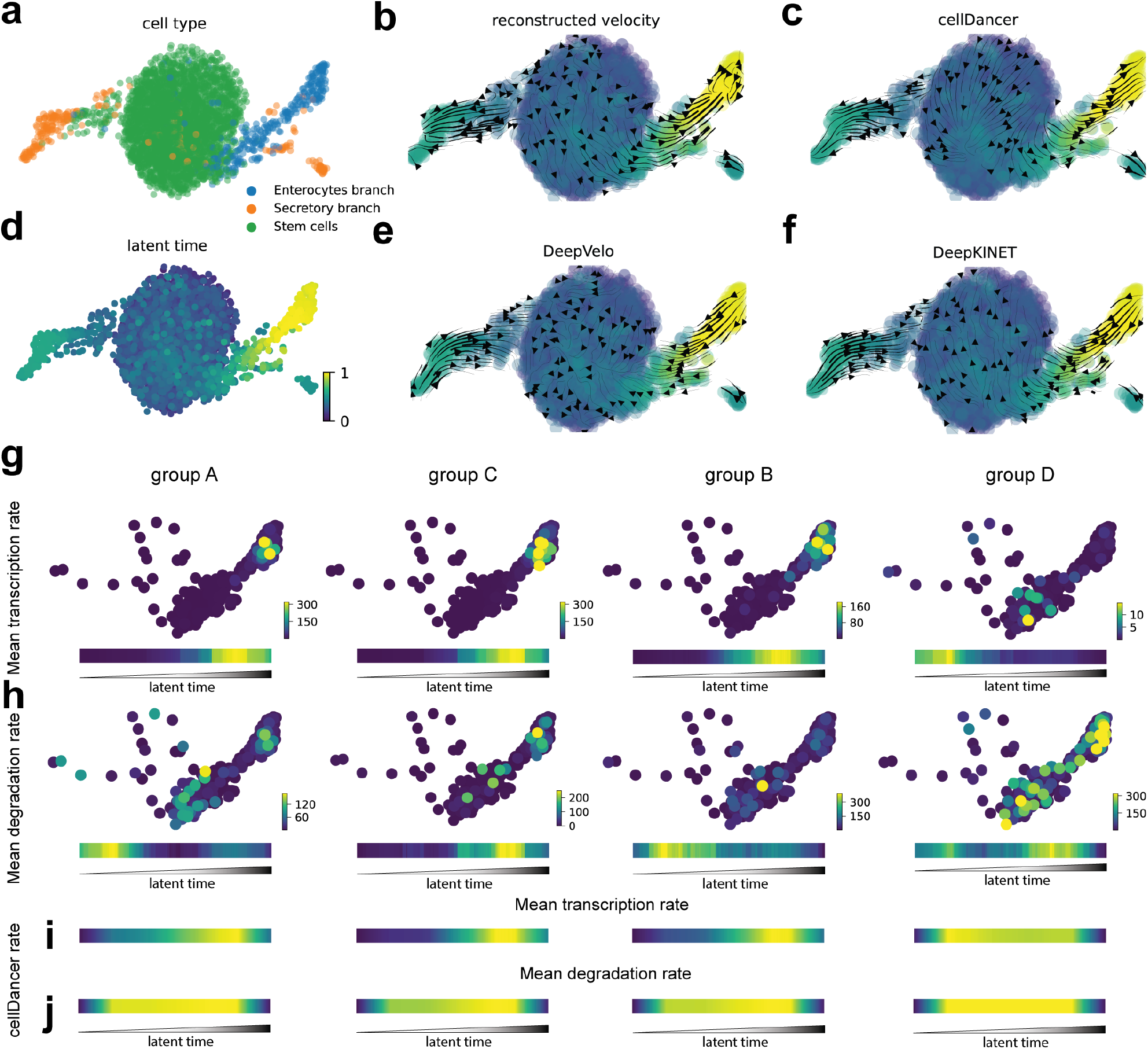
scRNAkinetics’ estimatation is congruent with kinetic rates of metabolic labeling scRNA-seq datasets. **a**, 2-D Umap visualization colored by cell branch. **b**, Reconstructed RNA velocity of scRNAkinetics based on the latent time derived from UnitVelo. **c**, RNA velocity estimated by cellDancer. **d**, Latent time estimated from UnitVelo. **e**, RNA velocity estimated by DeepVelo. **f**, RNA velocity estimated by DeepKINET. **g**, scRNAkinetics’ estimation of mean transcription rate for each group of genes in the enterocytes branch. The color bars on the right denote the display rate range. The heatmaps at the bottom show the relative mean transcription rate changes along the latent time. **h**, The mean degradation rate of each group of genes. The elements are the same as **g. i, j**, Mean transcription and degradation rate changes of each group of genes estimated by cellDancer. The 2D visualization is not displayed. The color bar indicates that cellDancer can not estimate the correct trends.

According to the original scEU-seq study, the researchers identified four groups of genes that exhibited intriguing gene regulation schemes in terms of different kinetics trends on the enterocytes branch. Generally, these genes can be divided into two categories, displaying either cooperative (group B and D) or destabilizing strategies (group A and C), respectively. The cooperative genes show a negative correlation between transcription and degradation, in contrast to the destabilizing genes, which exhibit a positive correlation between these processes. We examined the prediction of scRNAkinetics on these genes by plotting the mean transcription and degradation rate on the cells and displaying the relative rate changes along the latent time to illustrate the kinetic trends (Fig 3g,h). The results indicate that our method captures a weak destabilizing relationship for group A and a strong destabilizing relationship for group C. For groups B and D, scRNAkinetics predicts a weak and a strong cooperative relationship, respectively. Since cellDancer also predicts accurate velocity flow, we examined its predicted kinetic rates for these groups (Fig. 3i,j). However, cellDancer’s kinetic estimations do not exhibit the expected kinetic trends for these groups, indicating that the finer-grained kinetic estimation of cellDancer is still problematic.

### 2.4 scRNAkinetics leads to meaningful discoveries on conventional scRNA-seq datasets

Next, we applied scRNAkinetics on several real-world conventional scRNA-seq datasets to see whether the prediction was concordant with previous studies or not. First, we analyzed the erythroid gastrulation dataset, where a list of genes has been identified as transcription boosting genes. For this dataset, we incorporated cell differentiation potential estimated by cytoTRACE to inferring the kinetic parameters [22]. The reconstructed velocity correctly predicts the erythroid development direction (Fig. 4a). Nevertheless, the kinetic rates of *Hba-x* and *Hbb-y* display a transcription boosting behavior around the intersection between Blood progenitor 2 cells and Erythroid 1 cells (Fig. 4b,c). Besides this dataset, previous studies also showed that *HBB* and *KLF1* are likely to be the transcription boosting genes in the bone marrow dataset [25]. With scRNAkinetics, we calculated all the kinetic parameters and reconstructed the RNA velocity (Fig. 4d) which is aligned with our prior knowledge. We then investigated the kinetics rates of these two genes with the help of palantir pseudo-time derived by the stem cell prior [20]. The results indicate the transcription boosting of *HBB* is much stronger than that of the *KLF1* (Fig. 4e,f).

**Fig. 4.**
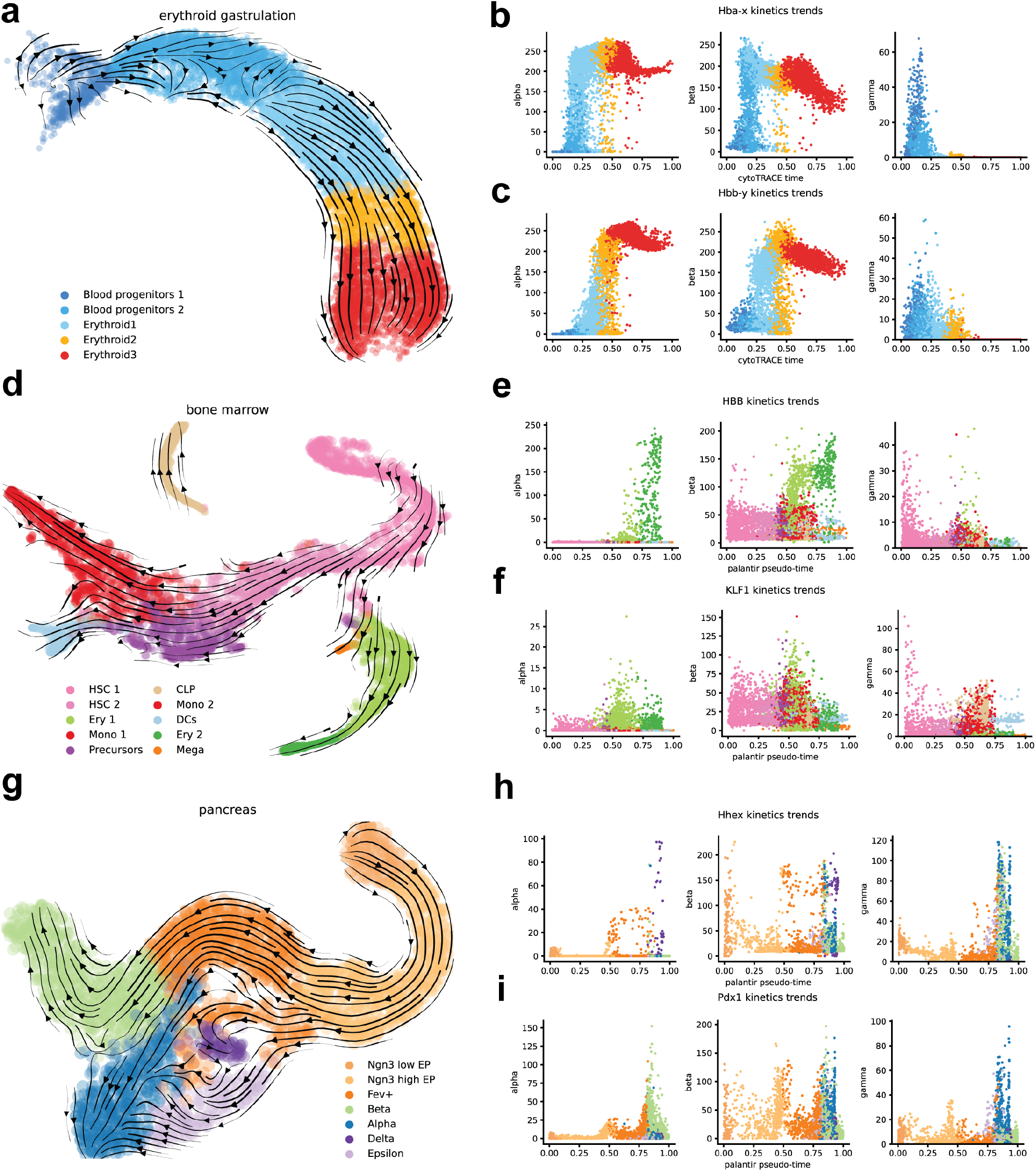
scRNAkinetics estimates reasonable kinetic rates on conventional scRNA-seq datasets. **a, d**, and **g** show the reconstructed velocity stream plot of scRNAkinetics for the three conventional scRNA-seq datasets. **b**,**c** Kinetics trends plot for transcription boosting genes in erythroid gastrulation dataset. **e**,**f** Kinetics trends plot for transcription boosting genes in bone marrow dataset. **h**,**i** Kinetics trends plot for delta cells driver gene (*Hhex*) and beta cells driver gene (*Pdx1*).

In addition to the transcription boosting genes, we are also interested in the kinetic rates of lineage determinant genes. Here, we introduced the well-studied pancreatic development dataset as an example. The kinetics analysis is based on the palantir pseudo-time derived by setting Ngn3 low EP cells as the stem cells. The reconstructed velocity correctly shows the differentiation direction but does not identify epsilon cells as a terminal state (Fig. 4g). This finding is different from previous CellRank analysis [28] but is congruent with previous experimental study [29], where researchers found epsilon cells still have the potential to differentiate into alpha cells. Moreover, we delved into the kinetic rates of *Pdx1* and *Hhex* genes which are beta cells and delta cells driver genes, respectively (Fig. 4h,i). The kinetics trends show these genes also display a similar transcription-boosting behavior in the corresponding cell type, indicating these genes are crucial to maintain cell differentiation.

### 2.5 scRNAkinetics validates the high correlation between transcription and splicing processes

In the cell nucleus, the processes of mRNA transcription and splicing are spatiotemporally coordinated by recruiting and assembling splicing factors along with the polymerase II elongation process [30,24]. Based on the fact that transcription and splicing are highly correlated, we hypothesized that our method should also observe this phenomenon by calculating the Pearson correlation between the estimated transcription rate and splicing rate for every gene. We then used a histogram to show the distribution of these correlations (Fig. 5a,b,c) for all genes in the three conventional scRNA-seq datasets. In contrast, we also displayed the Pearson correlation between transcription and degradation for these three datasets. The results suggest a highly correlated relationship pattern between transcription and splicing, while the pattern between transcription and degradation is inconsistent. This indicates that the underlying biological mechanisms of RNA dynamics can be captured by scRNAkinetics.

**Fig. 5.**
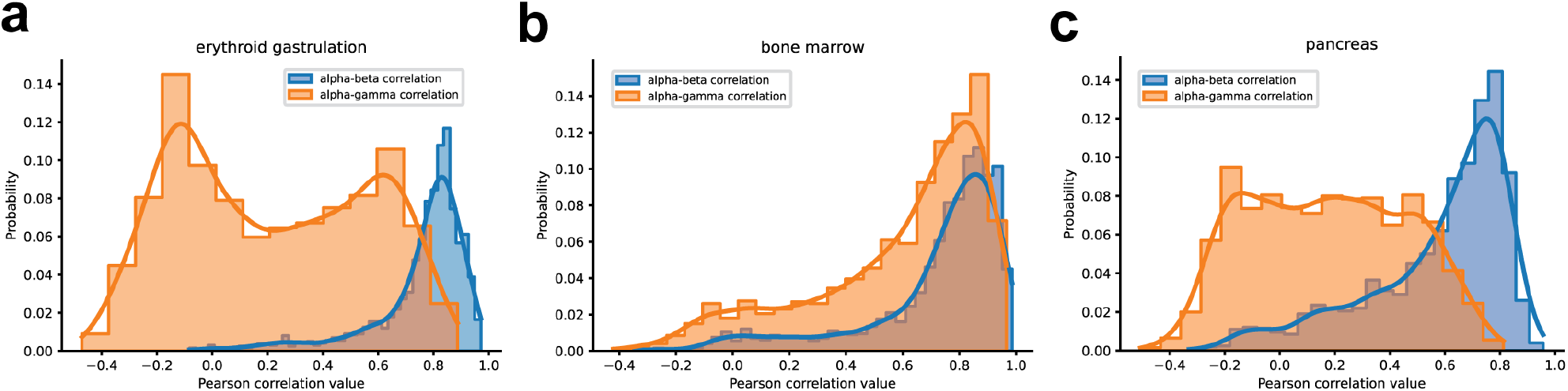
scRNAkinetics predicts transcription-splicing coordination. **a**, Histogram plots for transcription-splicing Pearson correlation and transcription-degradation Pearson correlation in erythroid gastrulation dataset. The Pearson correlation is calculated between each gene’s transcription rate, splicing rate, and degradation rate. Here we only show the correlation of transcription-splicing and transcription-degradation. **b**, Histogram plots for transcription-splicing and transcription-degradation Pearson correlation in bone marrow dataset. **c**, Histogram plots for transcription-splicing Pearson correlation and transcription-degradation Pearson correlation in pancreas dataset. These datasets show that most of genes have the high correlation between transcription and splicing and are consistent across different datasets.

## 3 Discussion

We developed scRNAkinetics as a new tool to infer single-cell RNA kinetics based on various biological priors. Unlike previous methods that use neural networks to predict kinetic rates, scRNAkinetics utilizes a rigorous numerical ODE solver to resolve the underlying kinetic rates for each cell. One major improvement over existing methods is the introduction of pseudo-time to solve the kinetics ODEs. Although this feature requires users to have more biological priors and understanding of the dataset, it is really worth it since scRNAkinetics can transform qualitative understanding of cell type transition into finer-grained kinetic parameters.

Validated on simulated and real-world datasets, scRNAkinetics is capable of handling the complex dynamic system in tissues and provides biologically meaningful results. However, we also noticed that scRNAkinetics still has some limitations. The first is that the method can only predict a relative kinetic rate instead of an absolute rate, which is due to the dependence on pseudo-time and the KNN smoothing procedure. The second limitation is the running time of scRNAkinetics, which is still slow compared to state-of-the-art single-cell methods; it can only estimate the kinetics of 1,200 samples in one hour. This could be further improved by optimizing parallelization and using compiled languages.

In summary, scRNAkinetics can be seen as a tool to transform pseudo-time into RNA velocity and kinetic parameters, and a tool to generalize the velocity estimation from a subset of genes [23] or latent space [31] to all genes in the original space. With the finer-grained kinetic information offered by scRNAkinetics, we can extend the current single-cell analysis framework from gene expression to kinetics analysis. Furthermore, implementing scRNAkinetics in numerical ODE solver prepares the framework for more complex RNA dynamic generalization in the multi-omics era. Considering all the facts about scRNAkinetics, we believe it will be helpful for researchers investigating single-cell kinetics *in vitro* and discovering new gene regulation schemes.

## 4 Methods

### 4.1 Datasets and preprocessing

In this work, we simulated or used several public available datasets to conduct experiments and analysis. The details are described below.

#### Simulated datasets

To assess the accuracy and limitations of our method, we utilized functions from previous packages to simulate the trajectories of the RNA dynamic ODEs with various settings [16,14]. According to the previous literature, current RNA velocity models are facing challenges when the steady state and constant kinetics rates assumption are violated [25]. These challenges include: 1. multiple-branches, where the phase plot shows two or more branches from multiple cell lineages with unknown evolving directions; 2. transcription boosting, where a gene’s transcription rate boosts under some differentiation regulations in the induction stage; 3. time-dependent kinetic rates, where the kinetic rates change along with time. Specifically, we used the cellDancer package to simulate the conventional, transcription boosting, and multi-branch forward scenarios and employed scVelo for the last scenario. For the conventional dynamics, parameters were set at *α* = 1, *β* = 1, *γ* = 1 during the induction stage and *α* = 0, *β* = 1, *γ* = 1 in the repression stage. In the transcription boosting stage, we increased the transcription rate (*α* = 3), while keeping other parameters constant. In the multi-branches forward scenario, the simulation of the second branch involved an elevated transcription rate (*α* = 2). Subsequently, the time-dependent degradation rate model was simulated using the scVelo package, adhering to the parameter settings of the original work: *α* = 2, *β* = 0.3 in the induction stage and *α* = 0, *β* = 0.3 in the repression stage. The degradation rate was time-dependent, defined as *γ* = 5 · (1 − *exp*(−0.3*t*)), applicable to both stages, with a maximum time set at 25.

In addition to simulating pure ordinary differential equation (ODE) trajectories across various models to evaluate the capabilities of our model, we introduced Gaussian noise into the datasets during the conventional induction stage to assess our model’s robustness. This noise was controlled using a scale factor, applied to the standard deviation of the Gaussian distribution. The noise (*ϵ*) was added according to the following formula:

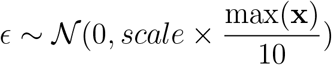

Here, **x** represents the vector of a specific gene’s abundance in either unspliced or spliced RNA, indicating that the noise level is relative to the maximum abundance of the RNA.

#### Real datasets

Besides proving our method’s correctness on the simulated datasets, we also incorporated several real scRNA-seq datasets from various tissues and technologies.

##### Metabolic labeling scRNA-seq datasets

The metabolic labeling data refers to using special chemical pounds to substitute the natural RNA base and then doing sequencing. By counting the number of labeled RNA after a time period, researchers can thus estimate the RNA transcription and degradation rate. In this work, we used the metabolic labeling organoid datasets from [6] to verify our method’s performance. This dataset contains four count matrices, “spliced unlabeled”, “spliced labeled”, “unspliced unlabeled”, and “unspliced labeled”. To perform the analysis in our framework, we summarized these four matrices as “unspliced” and “spliced” matrices by adding corresponding matrices together.

##### Conventional scRNA-seq datasets

To show our method’s capability on the widely-used conventional scRNA-seq datasets, we included three previously published datasets for further analysis. The first is the bone marrow datasets from [20] with labeled cell type and pseudo-time information. The second one is the murine pancreas developmental dataset at stage E15.5 from [32] with labeled cell type and palantir pseudo-time. The third one is the erythroid lineage of a mouse gastrulation dataset with only the cell type information from [33]. These datasets with preprocessed “unspliced” and “spliced” count matrices can be accessed from the original works or from the scVelo and CellRank datasets module.

#### Preprocessing

For simulated datasets, we directly use the data without further processing. For real datasets, we first employed our previously developed doublet filtering method to filter out doublets if necessary [34]. Then we followed the preprocessing steps in RNA velocity to filter out lowly expressed genes, normalize data, select highly variable genes (HVGs), and apply K-nearest neighbors (KNN) smoothing on both unspliced and spliced count matrices. To be specific, only genes with more than 20 counts in both matrices are kept, and then we selected 2000 HVGs and retained some interested genes for downstream analysis. For KNN smoothing, though the previous study has shown it affects the final results with different values [35], we followed the scVelo framework and set it as 30 for all real datasets.

### 4.2 scRNAkinetics framework

Generally, scRNAkinetics is a tool to decipher the underlying kinetic parameters for each cell with proper biological priors as guidance. Below is a detailed description of this method.

#### Pseudo-time derivation

As we mentioned before, the key problem in current RNA velocity/kinetics analysis is the lack of latent time guidance and solving underdetermined system. Hence, the first thing is to derive reliable pseudo-time as a guidance for the estimation of RNA kinetics in the next step. To do so, we employed biological priors to guide the pseudo-time estimation. Specifically, we summarize four kinds of priors (Fig 1) namely as stem cell prior, cell differentiation potency prior, RNA velocity prior, and cell lineage information prior. To incorporate these priors, there are several well-developed pseudo-temporal analysis methods. For the first prior, if we know the stem cell in the sequenced tissue, we can use palantir [20] or monocle [19] to set the stem cell and then get the pseudo-time estimation of all cells. For the second prior, SCENT [21] and cytoTRACE [22] utilize some hypotheses to estimate the differentiation potency and then transform potency into the pseudo-time value. For the third prior, velocyto [13], scVelo [14], and UnitVelo [27] can estimate RNA velocity with different mathematic model and transform the velocity into pseudo-time. For the cell lineage prior, PhyloVelo [23] can estimate velocity based on cell lineage information if available. Then the velocity can also be transformed into pseudo-time.

#### RNA dynamic model

Conventionally, RNA dynamics for each gene is described by a set of ODEs [8,13]:

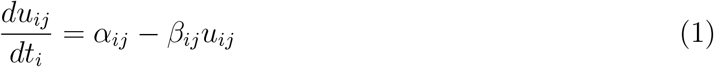

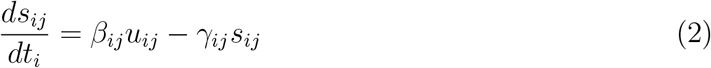

Here, *α*_*ij*_, *β*_*ij*_, and *γ*_*ij*_ represent the non-negative kinetic rates for transcription, splicing, and degradation of cell *i* and gene *j*, respectively. The variables *u*_*ij*_ and *s*_*ij*_ denote the abundance levels of the gene *j* of cell *i*, and *t*_*i*_ represents the real-time or pseudo-time, depending on the specific context of the analysis. Of note, this dynamic model can be extended to a more complex one when integrating single-cell multi-omics datasets, which may offer additional information, such as translation rate and transcription switch point [36,37].

#### Loss function

For a given group of cells **X** = {**x**_1_, **x**_2_, ⋯, **x**_*n*_} where **x**_*i*_ = {*u*_*i*,1_, *u*_*i*,2_, ⋯, *u*_*i,m*_, *s*_*i*,1_, *s*_*i*,2_, ⋯, *s*_*i,m*_} and pseudo-time points **t** = {*t*_1_, *t*_2_, ⋯, *t*_*n*_}. Without loss of generality, we assume *t*_1_ ≤ *t*_2_ ≤ ⋯ ≤ *t*_*n*_. Then, with **x**_1_ as the starting point, we use the DOPRI8 method [38,39] as the numerical ODE solver to predict the future states at **t**_*f*_ = {*t*_2_, ⋯, *t*_*n*_} based on the given RNA dynamic model. The future state is denoted as 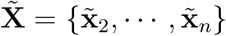. Of note, the predicted future states is actually a function of kinetic parameters, that is, 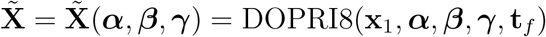. Here, ***α, β, γ*** are three vectors represent kinetic parameters for all *m* genes. To make the loss function can capture the small scale differences, we used the mean absolute error as the loss function:

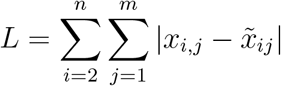

#### Optimization

In the optimization step, we used projected gradient descent to make sure the kinetic parameters are non-negative (Algorithm 1). All kinetic parameters are initialized with 1. We utilized the Adam optimizer [40] with a learning rate 0.1 and set the maximum iteration number as 2,000 throughout the paper. These parameters are chosen by making scRNAkinetics can faithfully recover the underlying kinetic parameters on the simulated datasets.

#### Implementation

For scRNAkinetics, we must determine the kinetic parameters for each cell and gene, which results in a complexity of *O*(*knm*) for datasets comprising *n* cells, *m* genes, and a *k*-nearest neighbors graph after the preprocessing step. To accelerate this process, we employed the vectorized JAX function for the concurrent estimation of all genes in a cell [41]. The preference for JAX is driven by its superior efficiency in the numerical integration of ODEs, compared to PyTorch-based ODE solvers, and it significantly outperforms the loop-based optimization approach in SciPy. Furthermore, we have integrated the multiprocess Python library, joblib, to parallelize computations across individual cells. With these enhancements, computing for a sample with *k* = 30 and *m* = 2000 is about 3 seconds on a single core of an Intel(R) Xeon(R) Silver 4216 processor.

## 5 Acknowledgements

## 6 Author contributions

Y.C., W.C., and H.H. conceived the project, S.F. processed the data, Y.C. conducted the experiments, Y.C. analyzed the results. K.N. and Z.H. reviewed the code. Y.C. wrote the manuscript and all authors reviewed the manuscript.

## 8 Tables

### Algorithm 1 Projected Gradient Descent

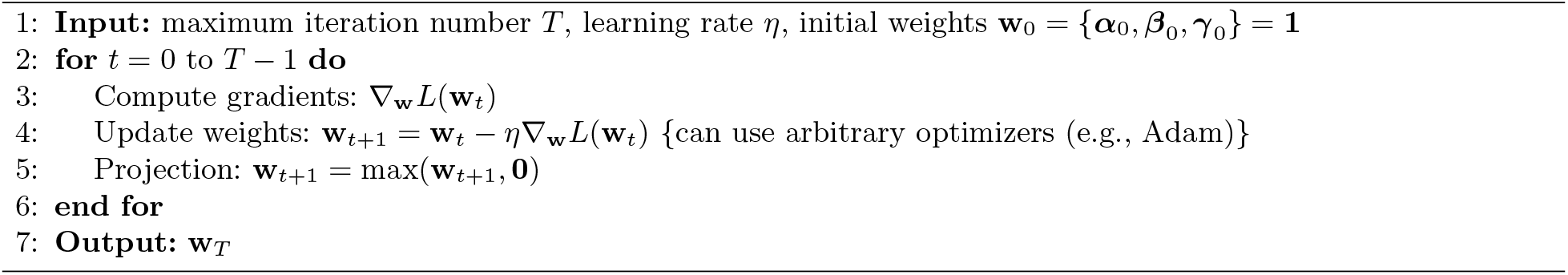

